# SL-Cloud: A Computational Resource to Support Synthetic Lethal Interaction Discovery

**DOI:** 10.1101/2021.09.18.459450

**Authors:** Bahar Tercan, Guangrong Qin, Taek-Kyun Kim, Boris Aguilar, Christopher J. Kemp, Nyasha Chambwe, Ilya Shmulevich

## Abstract

Synthetic lethal interactions (SLIs), genetic interactions in which the simultaneous inactivation of two genes leads to a lethal phenotype, are promising targets for therapeutic intervention in cancer, as exemplified by the recent success of PARP inhibitors in treating BRCA1/2-deficient tumors. We present SL-Cloud, an integrated resource and framework to facilitate the prediction of context-specific SLIs by using cloud-based technologies. This resource addresses two main challenges related to SLI inference: the need to wrangle and preprocess large multi-omic datasets and the multiple comparable prediction approaches available. We demonstrate the utility of this resource by using a set of DNA damage repair genes as the basis for predicting potential SLI partners, using multiple computational strategies. Context-specific synthetic lethality potential can also be compared using the framework. We demonstrate various use cases for our cloud-based computational resource and the utility of this approach for customizable and extensible computational inference of SLIs.

## Introduction

The concept of synthetic lethality (SL) refers to interactions between two genes in which loss of function of either gene alone does not impair cell viability, whereas inhibition of both genes is lethal (O’Neil et al., 2017). These interactions are attractive for designing cancer therapies, as targeting a gene whose synthetic lethal partner is permanently inactivated in cancer cells but exhibits wild-type expression in healthy cells should selectively kill cancer cells. The synthetic lethal interaction (SLI) between the poly (ADP-ribose) polymerase (*PARP*) genes and BRCA deficiency (functional loss of either B*RCA1* or *BRCA2*) is the first successful clinical application of the SL concept (Fong et al., 2009; Lord and Ashworth, 2017). Subsequent functional screens have proposed other synthetic lethal pairs, including the SWI/SNF chromatin remodeling complex members *SMARCA2*-*SMARCA4* (Hoffman et al., 2014) and *ARID1A-ARID1B* (Helming et al., 2014), as well as the Werner syndrome RecQ-like helicase (*WRN*) gene in *MYC* overexpressing cancers (Moser et al., 2012) and microsatellite unstable cancers (Chan et al., 2019; Kategaya et al., 2019; Lieb et al., 2019). Although SL-based therapeutics are promising, other drugs for clinical use designed using an SL-based rationale are still under development. There is, therefore, a continued need to discover additional synthetic lethal gene pairs and to develop automated methods that use various data types to predict clinically relevant synthetic lethal pairs that can be nominated for further testing and therapeutic development (Huang et al., 2020).

Functional screening using siRNA/shRNA technology or, more recently, CRISPR-based targeting libraries is a leading method of SLI discovery (O’Neil et al., 2017). However, identifying robust synthetic lethal gene pairs is challenging, in part due to biological factors such as tumor heterogeneity and incomplete penetrance, i.e. context-dependent SL (Chan et al., 2010; Henkel et al., 2019; Nijman and Friend, 2013; Ryan et al., 2018). To complement functional screening efforts, multiple computational prediction strategies have been pursued (reviewed by (O’Neil et al., 2017)). Early approaches inferred SLIs in humans via orthology mapping based on genetic interaction networks from experimentally-tractable model organisms such as *Saccharomyces Cerevisiae* (Conde-Pueyo et al., 2009; Kirzinger et al., 2019; Srivas et al., 2016) and *Mus Musculus* (Gurley and Kemp, 2001). Alternative strategies rely on the integrated analysis of multi-omic profiling and functional screening of patient-derived or cancer cell line based datasets to predict SLIs. Briefly, these approaches use statistical and/or heuristic methods, such as implicating SL gene pairs via mutually exclusive loss-of-function mutations, shared pathways or protein complex membership (Das et al., 2019; Jerby-Arnon et al., 2014; Lee et al., 2018; Liany et al., 2020; Wappett et al., 2016; Ye et al., 2016). Furthermore, to facilitate interactive exploration of predicted SLIs, several web portals or SLI databases have been published, such as Syn-Lethality (Li et al., 2014), SynLethDB (Guo et al., 2016), the Synthetic Lethality BioDiscovery Portal (SL-BiodP) (Deng et al., 2019), the Cancer Genetic Interaction Database (CGIdb) (Han et al., 2019), and, more recently, SynLeGG (Wappett et al., 2021). These tools present pre-computed synthetic lethal pairs based on the most comprehensive datasets available at the time of publication. This necessarily excludes potential SLIs discoverable by either algorithmic advances or developments in functional screening technologies in terms of scope and throughput. Additionally, there is limited flexibility to explore the existing set of putative SLIs or to change any parameters in the prediction algorithms to better understand how the SL inference was made.

In addition to the aforementioned limitations, benchmarking the performance of different prediction approaches is difficult, as no *bona fide* set of gold-standard SL interactions exists. Because of the complexity of these prediction approaches and managing the amount of data on which predictions are based, a framework to assess and compare multiple prediction approaches simultaneously on the same dataset does not exist. Many of the current computational approaches for SLI prediction do not provide customized scripts or a facile connection to large public data resources, making it difficult to use the wealth of publicly available data that can be repurposed for SLI prediction. These reasons also account for the multiplicity of SL databases, with the accompanying challenge presented to the user of having to select the most appropriate database and tools for their needs.

To address the challenges associated with computational prediction of SL, we developed Synthetic Lethality Cloud (SL-Cloud), a cloud-based computational resource to support SL investigations. We mined large-scale publicly available multi-omics and functional screening datasets such as The Cancer Genome Atlas (TCGA) (Hutter and Zenklusen, 2018), the Cancer Cell Line Encyclopedia (CCLE) and the Cancer Dependency Map (DepMap) (Dempster et al., 2019; Ghandi et al., 2019; McFarland et al., 2018; Meyers et al., 2017) to infer SL. To eliminate the need for individual investigators to re-download these large multi-omic datasets, we make use of the cloud-hosted Google BigQuery tables provided by the Institute for Systems Biology Cancer Gateway in the Cloud (ISB-CGC) (Reynolds et al., 2017). Additionally, we have collected, reformatted, and structured other large-scale datasets not otherwise available on the ISB-CGC into Google BigQuery (cloud-enabled SQL-queryable data warehouse) tables for use in developing SL inference workflows. We implemented three SL inference pipelines, including the Conserved Genetic Interaction (CGI) workflow, an orthology-mapping approach based on SLIs discovered in *S. Cerevisiae* (Srivas et al., 2016); a mutation-dependent synthetic lethality prediction (MDSLP) workflow that combines mutation and essentiality screening data to infer SL pairs from cancer cell line data; and a re-implementation of the previously published data-mining synthetic lethality identification pipeline (DAISY) (Jerby-Arnon et al., 2014). These workflows are encoded in Jupyter notebooks that use functions from the provided Python scripts and embedded SQL queries that pull down relevant data from the cloud-based BigQuery tables for SL prediction. SL-Cloud brings together computational pipelines alongside large-scale datasets via cloud infrastructure to facilitate highly scalable and customizable analyses.

## Materials and Methods

### Data Collection

We identified datasets hosted on the ISB-CGC platform that are relevant for SL inference, e.g., TCGA Pan-Cancer Atlas gene expression data. and designated them for use in this project (see Table 1 for details). Additional datasets, particularly those from the Cancer Dependency Map (DepMap) including genomic characterization data (on mutations, copy number, and gene expression) from the CCLE project (Ghandi et al., 2019), gene effects estimated from CERES for CRISPR-Cas9 essentiality screening from Project Achilles (Meyers et al., 2017), and cancer cell line gene dependency scores estimated from DEMETER2 v6 from three large RNAi screening datasets (McFarland et al., 2018) were downloaded from the DepMap data portal. Similarly, comprehensive synthetic lethal interaction data from synthetic genetic array (SGA) perturbation screens in S. Cerevisiae were downloaded from TheCellMap resources. Human-to-S. Cerevisiae orthology mapping information for this paper were retrieved from the Alliance of Genome Resources (Release 3.0.1). All datasets not previously available via the ISB-CGC were collated in a Google Cloud Project (GCP Project ID: “syntheticlethality”) established for the current project. All datasets were downloaded, parsed, and stored as BigQuery SQL-like tables. We downloaded and stored gene metadata tables as BigQuery tables to facilitate table integration across datasets based on the “Entrez Gene ID” column and for maintaining HGNC gene symbol compatibility. All of these datasets are accessible programmatically via SQL-like queries through the Google Cloud API for various languages, e.g., R, Python, Ruby, Java, etc., and through a web-based interface.

### Data and Code Availability

All datasets supporting the current study are hosted on the ISB-CGC (Reynolds et al., 2017) in existing Google BigQuery SQL database tables (Table 1). Additional publicly available datasets relevant to SL inference were downloaded from the sources outlined in Table 1 and were uploaded to new BigQuery tables in the Google Cloud Project (GCP) for this paper (GCP Name: “syntheticlethality”). Project code and documentation describing how to access and use this resource are available on the project github page at https://github.com/IlyaLab/SL-Cloud.

### Reimplementation of Synthetic Lethal Prediction Algorithms

#### Data-mining synthetic lethality identification pipeline

The data-mining synthetic lethality identification pipeline (DAISY) was implemented as described previously (Jerby-Arnon et al., 2014). DAISY uses three inference modules: pairwise gene co-expression, survival of the fittest, and functional examination to enable the detection of both synthetic lethal and synthetic dosage lethal gene pairs.

In the pairwise co-expression module, DAISY makes inferences based on the assumption that synthetic lethal gene pairs play a role in related biological processes and are co-expressed. Gene expression is measured for patient-derived data from TCGA and cancer cell line-derived data from CCLE. Pairwise co-expression is estimated from the Spearman correlation which we calculated between each gene of interest (each item in the query gene list) and all other genes. Candidate synthetic lethal gene pairs are those with correlation coefficient greater than 0.5 and whose Bonferroni-corrected *P* value was smaller than 0.05. We used SQL query-based implementations of the Spearman correlation and Wilcoxon test calculations to estimate effect sizes and *P* values when retrieving the data, because that approach is faster than using the standard Python libraries.

The other two inference procedures are built on the definition of SLIs in that functional loss of both genes in a synthetic lethal pair decreases cell viability and is, therefore, selected against. The genomic survival of the fittest inference module is based on the statistical test of the copy number alteration of the gene in the search domain, given whether or not the gene of interest is inactive (overactive) or not. The gene of interest in a sample is considered inactive if its expression is less than 10th percentile across all samples and its copy number alteration is less than -0.3 or if it has a nonsense, frame shift or frame-del mutation. The gene of interest in a sample is considered *overactive* if it has gene expression exceeding the 90th percentile across all samples and its copy number alteration is greater than 0.3 (over-activity is used in synthetic dosage lethal pair prediction). The one-sided Wilcoxon rank-sum (Mann-Whitney U) test was applied to the copy number alteration of the candidate synthetic lethal pair of each gene of interest. The higher copy number of the candidate synthetic lethal pair for the samples whose gene of interest is inactive (overactive) is considered as an indicator of the genes being in a synthetic lethal or synthetic dosage lethal relationship. The SL/SDL pairs whose Bonferroni - corrected p-value is less than 0.05 were returned. This inference procedure was applied on Pancancer Atlas and CCLE data separately. The union of results from PanCancer Atlas and CCLE was used.

The rationale for the functional examination inference module is that if the synthetic lethal partner of a gene is inactive in a given sample, subsequent inactivation of that gene will be lethal. Therefore, for a gene of interest, we first defined two groups for the test, one in which the gene was inactive and the other in which it was not. We then performed a one-sided Wilcoxon rank-sum (Mann-Whitney U) test on the knockdown/knockout sensitivity of candidate synthetic lethal pairs of interest. Lower viability that is associated with higher knockout/knockdown sensitivity is an indicator of a potential SLI. The synthetic lethal pairs for whom the test result *P* value was lower than 0.05 were returned. This inference procedure was applied to the gene-dependency scores or gene effect scores for the shRNA and CRISPR datasets separately. We reported the union of results from shRNA and CRISPR-based datasets.

The synthetic lethal pairs and their corresponding *P* values were listed for each data source and DAISY module. In addition, for all DAISY modules, we applied statistical testing where an appropriate number of samples with particular features was available (typically n >= 5). For example, for the SoF module, we applied the test only where the number of inactive/overactive samples was greater than five and the total number of samples was greater than 20. The thresholds for the gene inactivation/overactivation decision, *P* value, and mutation type can also be adjusted by users for their own analyses, and inclusion/exclusion of specific mutation classes to decide whether a gene is inactive is optional. Users can also use different multiple-test correction approaches. The Spearman correlation coefficient and *P* value thresholds and the method used for multiple-test correction are parametrized, and the user can set customize their selection of thresholds and multiple-test correction methods from among those defined in the ‘statsmodels.stats.multitest.multipletests’ Python library function.

#### Mutation-dependent synthetic lethality prediction

The MDSLP is based on genetic variants, gene-dependency scores, or gene effects from the cancer cell line data in the DepMap Portal (see sources in Table S1). The genomic variants data from the CCLE project, gene-dependency scores estimated from CERES for CRISPR-Cas9 essentiality screening from project Achilles (Meyers et al., 2017) and gene-dependency scores estimated from DEMETER2 from three large RNAi screening datasets (McFarland et al., 2018) were used. We developed a pipeline that integrates the genetic information and the functional screening data to evaluate mutation-based conditional dependence, using either the CRISPR or the shRNA dataset. For this pipeline, tumor types can be selected by the users. The variants selected for this pipeline include those that alter the amino acid in the protein structures or protein expression which may further impact the function of the gene product. These alterations include Splice_Site, Frame_Shift_Del, Frame_Shift_Ins, Nonstop_Mutation, In_Frame_Del, In_Frame_Ins, Missense_Mutation, Nonsense_Mutation, Nonstop_Mutation, Start_Codon_Del, Start_Codon_Ins, Start_Codon_SNP, Stop_Codon_Del, Stop_Codon_Del, Stop_Codon_Ins, and De_novo_Start_OutOfFrame.

For the CRISPR data-based pipeline, we used CCLE mutation, Achilles gene effect, and sample_info data from DepMap (version 20Q3). After selecting tumor types, or the pan-cancer analysis option, for each selected mutated gene, we grouped the cell lines into either the mutated or the wild-type group, then tested whether the knockout effects or the gene dependency scores for the two groups show statistically significant differences using a t-test, followed by Benjamini-Hochberg (BH) adjustment. Effect size (Function 1) was used to measure the difference between the two groups. For each measurement, only the sample size for each group larger than five was considered. For the shRNA data-based pipeline, cancer cell line gene dependency scores derived from DEMETER2 (version 6) from a combined dataset of Achilles (Meyers et al., 2017), DRIVE (McDonald et al., 2017), and shRNA screen in breast cancer cell lines (Marcotte et al., 2016) were used. The mutation data and sample annotation were for the DepMap 20Q3 dataset. Significant differences are defined for gene pairs with BH-adjusted P value smaller than 0.05. For the validation of the well-known synthetic lethal gene pairs, BH-adjusted P values smaller than 0.1 were also considered to indicate significance. Significant gene pairs with effect size smaller than 0 are predicted to be SLIs.

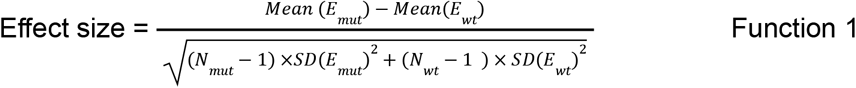

where *Emut* and *Ewt* represent the vector of knockout effects (or dependency scores) in the mutated and the wild-type groups, respectively. *Nmut* and *Nwt* are the numbers of samples in the mutated and the wild-type groups, respectively. SD is the standard deviation.

#### Conserved genetic interaction (CGI) workflow

The basis of this pipeline is the identification of high-confidence SLIs from digenic knock-out screens performed in *S. Cerevisiae*. Costanzo et al. measured double or single mutant colony fitness using synthetic genetic arrays (Costanzo et al. 2016). Genetic interactions are inferred by comparing double to single mutant colony growth as a measure of organismal fitness due to gene essentiality. If the cell viability of a double mutant colony is higher or lower than that of two single mutants then positive or negative genetic interactions are inferred using a quantitative fitness metric or generic interaction score. Synthetic lethal interactions are defined as the genetic interactions with low negative scores (< -0.35) at the extreme end of the interaction score distribution. Leveraging this dataset, we mapped yeast to human genes using yeast-human orthology information to identify presumed conserved human synthetic lethal pairs.

#### Pathway analysis of synthetic lethal partners of DNA damage repair genes

We downloaded a community-curated set of DNA damage repair (DDR) genes from (Knijnenburg et al., 2018) and evaluated the synthetic lethal gene pairs for each gene from this gene set with all available pipelines. To select synthetic lethal gene pairs corresponding to the DDR genes, we chose the predicted pairs with FDR < 0.05. The resulting list of synthetic lethal gene pairs was subjected to pathway and gene ontology enrichment analysis using the g:Profiler R Bioconductor package (Raudvere et al., 2019). Significant pathways or gene ontology biological processes (GOBPs) were identified (*P* < 0.05; intersection size > 5). The redundant GOBPs were reduced using REVIGO (Supek et al., 2011) based on the hierarchy of GO terms and clustering analysis based on semantic similarity.

#### Tumor type specific SL analysis

Tumor type-specific SL analysis is possible with the MDSLP pipeline. We used *ARID1A* as an example, ran the MDSLP-CRISPR and MDSLP-shRNA pipeline, and obtained the potential synthetic lethal partner genes for *ARID1A* by estimating the significance of the difference in the gene knockout or knockdown effects when comparing the cell lines in the *ARID1A* wild-type group and the *ARID1A*-mutated group for one specific tumor type. Student’s t-test followed by BH adjustment was used to estimate the significance. Only tumor types with at least five cell lines with both *ARID1A* mutations (functional mutations same as those described in the “Mutation-dependent synthetic lethality prediction” section) and gene knockdown or knockout effects data are taken into consideration for this analysis. Significance is considered as BH adjusted p-value smaller than 0.05.

The DAISY pipeline also enables tumor type-specific analysis. We manually added TCGA subtypes according to the tumor-type annotation in the CCLE datasets. The TCGA Pan-Cancer Atlas and CCLE samples were filtered based on the TCGA subtypes that are available in the CCLE project, and the DAISY algorithm was applied to this subset of data, using the same settings/thresholds as in the global analysis. For the functional examination and survival of fittests modules, the analysis was performed only if the number of inactive samples was greater than 5 and the overall sample size was greater than 20 per dataset individually. The pairwise co-expression module was performed if the overall sample size was greater than 20.

## Results

### SL-Cloud

We present an overview of the SL-Cloud framework in Figure 1. This resource includes pre-processed publicly available cancer genomics datasets hosted on a cloud resource platform and a set of SL inference workflows implemented as Python scripts. Our framework collates commonly used public data resources relevant for SL inference and combines them with analysis workflows that demonstrate how to infer SLIs by leveraging the cloud-based resources stored on the ISB-CGC platform. These components represent an ecosystem that integrates software and data to enable the large-scale prediction of SLIs from existing cloud-hosted datasets. In the following sections, we briefly describe the publicly available datasets that we identified as relevant for SL inference and we describe examples of use cases in which we re-implemented published SL inference workflows using these cloud-stored datasets.

**Figure 1.**
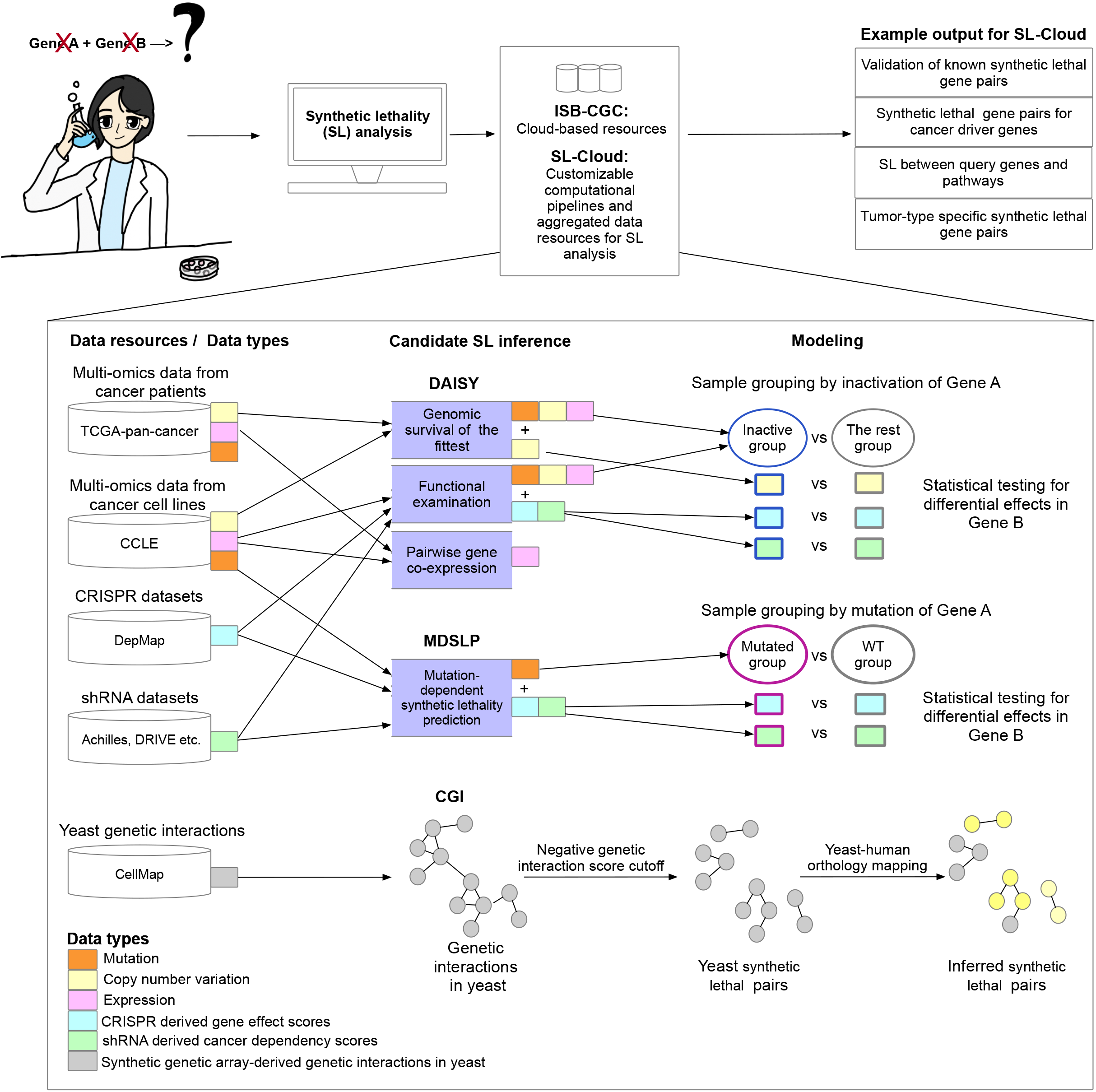
Overview of the Synthetic Lethality Cloud (SL-Cloud) framework. Schematic overview of the computational pipelines and data resources aggregated in this framework to facilitate investigation of synthetic lethal interactions. Users with specific research questions (top-left) reuse inference pipelines from the provided scientific computing notebooks to query public cancer genomics datasets from the vast resources provided by the ISB-CGC and additional datasets pre-processed in SL-Cloud. Three candidate synthetic lethal pair inference pipelines were implemented, including the data mining synthetic lethality identification pipeline (DAISY), mutation-dependent synthetic lethality prediction (MDSLP), and conserved genetic interaction (CGI). Different data resources and data types were used for different inference pipelines. For example, the DAISY and MDSLP pipelines rely on statistical testing over the multiple omics data and functional screening data such as CRISPR and shRNA datasets for human cancers, whereas the CGI pipeline is based on an ortholog mapping of the SL interactions identified in yeast.

### Large-scale datasets relevant for SL prediction

We have developed an aggregated resource that takes advantage of democratized access to public cancer genomics datasets through the National Cancer Institute’s Cancer Research Data Commons. Specifically, we use existing data hosted by the ISB-CGC and stored as Structured Query Language (SQL)-like database tables using Google BigQuery technology (Reynolds et al., 2017). Example datasets include TCGA patient-level data on somatic mutations, gene expression and copy number variation across 33 cancer types (Figure 1)(for details see Table S1, which includes the URLs for specific resources). We identified additional data resources that were pertinent for SL inference but were not yet available on the ISB-CGC platform. These datasets include genetic interaction datasets derived from model organism interaction screens, such as TheCellMap (Costanzo et al., 2016), and from human pan-cancer cell line molecular characterization and functional screening datasets, primarily from the DepMap initiative (Dempster et al., 2019; Ghandi et al., 2019; McFarland et al., 2018; Meyers et al., 2017). Building on the infrastructure of the ISB-CGC, we established a new SL-focused cloud resource project that incorporates resources not already present on the ISB-CGC (Google Cloud Project: “syntheticlethality”, Table S1). This will enable researchers to upload private datasets and analyze them alongside the wealth of public data.

### Synthetic lethal interaction prediction workflows

We re-implemented published SLI inference workflows and redistributed them as a set of Python scripts embedded in a set of Jupyter notebooks with the appropriate documentation describing the methodology. These notebooks offer code optimization and integration with the ISB-CGC through the BigQuery interface to access the relevant pre-processed large-scale cancer genomics datasets described above. The resource facilitates custom analyses for particular research questions or disease contexts. Additionally, this framework provides extensibility by virtue of the modular design of the base framework shown in Figure 1. End-users can combine high-quality public data with their own laboratory-generated data to perform integrated analyses more easily without the need to download and pre-process large-scale public cancer genomics data.

We implemented three workflows to demonstrate the utility and flexibility of this framework for SL prediction. First, we re-implemented the published workflow, DAISY (Jerby-Arnon et al., 2014), using up-to-date, large-scale data resources as described above (Figure 1; Table S1). DAISY applies multiple inference procedures that include genomic survival of the fittest (SoF), the detection of infrequently co-inactivated gene pairs by using somatic mutation, copy number alteration and gene expression data; functional examination (FunEx), the identification of gene pairs in which inactivation or over-activation of one gene induces essentiality of a partner gene, using functional screening data; and pairwise gene co-expression, the detection of significantly correlated gene pairs, thereby implicating genes in related biological functions. Inference of synthetic dosage lethality (SDL), whereby overactivation of one gene causes its interaction partner to become essential, is also implemented in DAISY. DAISY SL predictions are gene pairs that are found by all three inference modules. Each individual module can also provide evidence of SL potential independently. The pipeline we have implemented enables users to list synthetic lethal pairs from each pipeline, for each dataset, and aggregate them or use them independently. The DAISY pipeline enables users to perform pan-cancer or tissue type-specific analyses.

We also implemented a mutation-dependent synthetic lethality prediction (MDSLP) workflow based on the rationale that, for tumors with mutations that have an impact on protein expression or structure (functional mutation), the knockout effects or inhibition of a partner target gene show conditional dependence for the mutated molecular entities (Figure 1). Leveraging the public cancer cell line datasets including gene mutation data from CCLE, and functional screening data generated by either shRNA or CRISPR technology from DepMap (Dempster et al., 2019; Ghandi et al., 2019; McFarland et al., 2018; Meyers et al., 2017), we integrated these data modalities to evaluate mutation-based conditional dependence. This pipeline enables users to test statistically whether the knockout or knockdown effects for one gene will be altered if another gene is mutated in specific contexts, such as in pan-cancer, or tumor type-specific cell lines. The increase in gene knockout or knockdown sensitivity provides evidence to support potential SLIs.

Finally, we present a workflow that leverages cross-species conservation to infer experimentally derived SLIs in yeast to predict relevant synthetic lethal pairs in humans. We implemented the conserved genetic interaction (CGI) workflow based on published methods described in (Srivas et al., 2016). In SL-Cloud, we downloaded and preprocessed TheCellMap dataset, the most comprehensive *S. Cerevisiae* genetic interaction network inferred from synthetic genetic array (SGA) screens from (Costanzo et al., 2016) (see details in Table S1). Genetic interactions are inferred if the combined effect of a double mutant on cell viability differs from that of the combination of single mutant effects. SLIs are defined as negative genetic interactions in which the cell viability of a double-mutant colony is lower than that of the respective single-mutant colonies. We provide the inferred SLIs in humans by yeast-to-human ortholog mapping.

### Use cases

To demonstrate the utility of SL-Cloud, we will present representative examples of the type of SL inference and analysis enabled by this cloud-based platform and highlight opportunities for performing SL discovery.

#### *In silico* validation of known or suspected SLIs

Synthetic lethality between *BRCA* deficiency and *PARP1/2* is well documented and is the rationale behind the design of PARP inhibitors such as olaparib, rucaparib, and niraparib (Ashworth and Lord, 2018). These agents are approved for treating *BRCA*-mutated ovarian cancer and advanced breast cancer. To perform an *in silico* validation analysis of this well-established SLI, we applied MDSLP to gene mutation and essentiality screen data from pan-cancer cell lines (Dempster et al., 2019; Ghandi et al., 2019; McFarland et al., 2018; Meyers et al., 2017). As shown by the MDSLP-shRNA workflow and consistent with our expectations, functional mutations of *BRCA2* showed significant sensitivity to *PARP1* knockdown (two-sided t-test, *P* < 0.01, FDR < 0.1) (Figure 2A). We applied the same pipeline to gene essentiality data derived from CRISPR screens, but did not find this expected interaction. The MDSLP workflow using CRISPR-derived datasets revealed that the functional mutation of *BRCA2* shows a synthetic lethal partnership with *PARP2* (*P* < 0.01, FDR < 0.05). shRNA-derived and CRISPR-derived *BRCA2* synthetic lethal pairs showed limited overlap (Figure 2B). Only 6.6% (48 out of 729) of the BRCA2-related synthetic lethal pairs nominated from shRNA-derived inference with threshold of FDR < 0.1 were also predicted using CRISPR essentiality screens. From among the 1433 potential partner genes predicted by any of the resources, only 48 partner genes were predicted by two resources (Figure 2A). Of these, *WRN, TSC2, RPL22L1* showed the most significance with both CRISPR-derived and shRNA-derived inference (FDR < 0.01 for both inference procedures).

**Figure 2.**
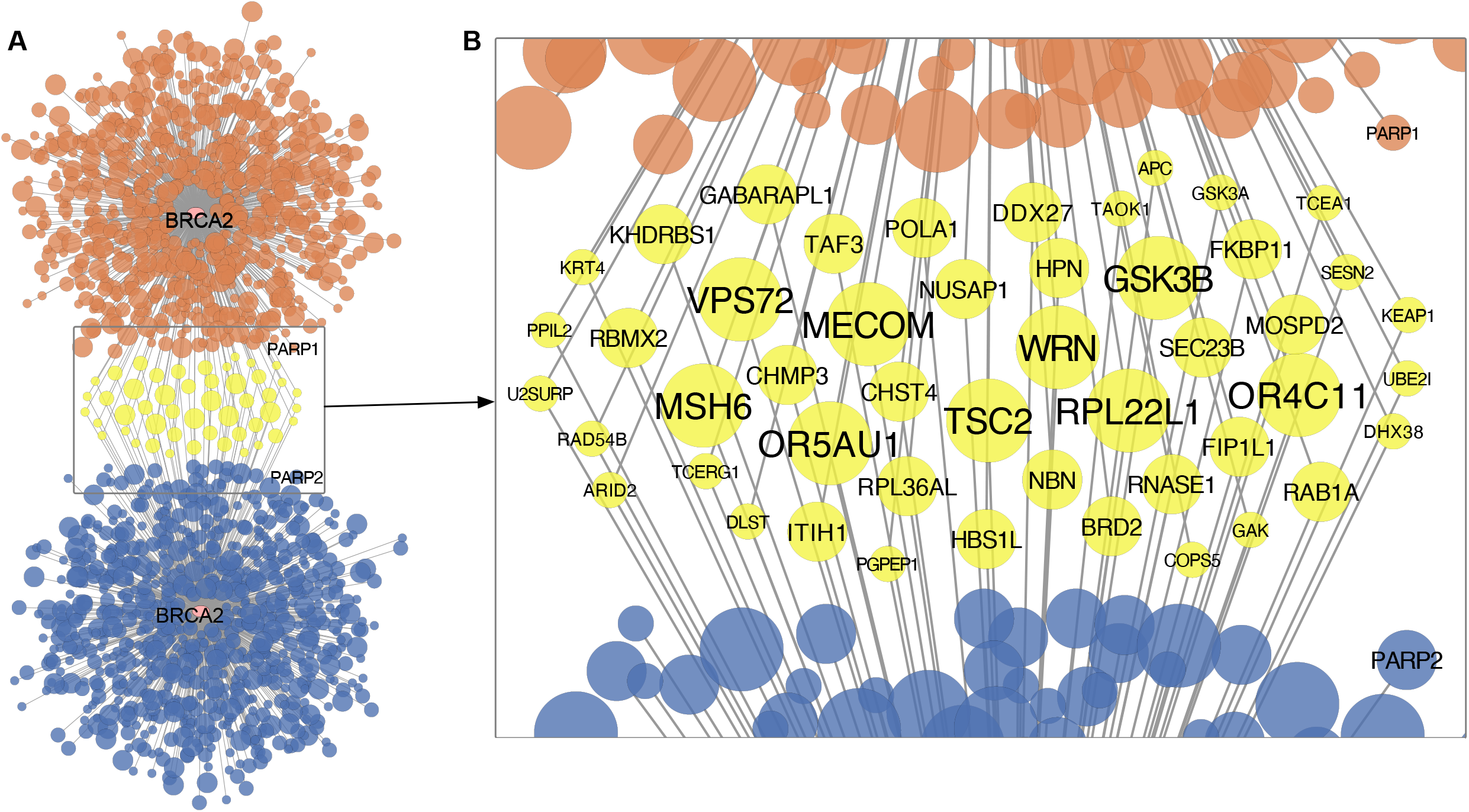
Predicted synthetic lethal interaction partners of *BRCA2* based on cancer cell line dependency datasets. **A**. Network-based representation of predictions generated by the mutation-based conditional dependency pipeline using either shRNA (upper panel, orange) or CRISPR (lower panel, blue) functional screening datasets. **B**. Potential synthetic lethal partners of *BRCA2* as predicted by both the CRISPR and shRNA functional screening data. Each node represents a gene, and edges represent potential synthetic lethal interactions. The node color encodes overlap between the gene-dependency assay type, with yellow representing synthetic lethal pairs detected in both the CRISPR data and the shRNA functional screening data. The node size and font size indicate the strength of the statistical relationship, with high-confidence pairs having larger node sizes or font sizes. FDR levels: 0.01 (largest), 0.05 (median), and 0.1 (smallest).

Interestingly, we did not predict any *BRCA2-*related synthetic lethal pairs from the other two workflows implemented in SL-Cloud. *BRCA2* has no yeast homolog and, therefore, conserved interactions could not be inferred by the CGI pipeline. DAISY nominated no synthetic lethal partners for *BRCA2* with its default settings but predicted potential *BRCA1-PARP1/2* SLIs across all three of its component inference modules with non-default parameter settings. Both gene pairs, *BRCA1-PARP1* and *BRCA1-PARP2*, showed statistically significant co-expression, with their correlation coefficients ranging from 0.26 to 0.59 across patient-derived and cancer cell line datasets [Figures S1A(i,ii) and S1B(i,ii)] (*P* < 0.01). In addition, we found statistical support for a *BRCA1* and *PARP1/2* SLIs by the SoF inference procedure [Figures S1A(iii) and S1B(iii,iv)] (*P* < 0.05), whereas the FunEx module found statistical support for a *BRCA1-PARP1* SLI [Figure S1A(iv)] (*P* < 0.01), but not for a *BRCA1-PARP2* SLI, based on the cancer cell line gene-dependency CRISPR or shRNA datasets. In summary, the *BRCA1-PARP1* interaction is supported by all three DAISY inference modules, whereas only two modules support the *BRCA1-PARP2* interaction.

This example demonstrates how a platform such as SL-Cloud can easily facilitate the exploration of the SLIs for a particular gene by using orthogonal SLI prediction workflows and multiple datasets to assess the stability (relative to algorithmic settings and parameters) and reproducibility of particular SLIs of interest. For the established SLI between BRCA deficiency and PARP1/2 enzymes, we see variation in the output of multiple prediction approaches and datasets in confirming this *bona fide* SLI. These analyses highlight some of the challenges related to SL prediction, including unaccounted-for variation resulting from differences in the datasets used to make the SL prediction and the implicit or explicit assumptions made by the underlying analytical approaches. These findings inform a better understanding of the mechanisms driving SLIs and inform algorithmic development for improving SLI prediction.

#### Pathway-based SL analysis

SLI partners tend to form functional interaction networks (Costanzo et al., 2016; Jerby-Arnon et al., 2014). For example, Ku et al. report that synthetic lethal screen hits are reportedly more robust at the pathway rather than at the gene level (Ku et al., 2020). To demonstrate pathway-based SLI analysis, we analyzed SLI-related genes in the DNA damage and repair (DDR) pathway. DDR deficiency due to loss-of-function alterations by mutation, deletion, or epigenetic silencing is prevalent across lineages affecting approximately 33% of all cancers in TCGA (Knijnenburg et al., 2018). Impaired DDR leads to genomic instability, and tumors exhibiting DDR loss are prone to DNA-damaging agents and, therefore, potentially vulnerable to inhibitors that target compensatory DDR pathways via a synthetic lethal mechanism (Lord and Ashworth, 2012). Using a well-curated set of 276 genes annotated for involvement in DNA damage repair from (Knijnenburg et al., 2018) we predicted synthetic lethal partners from the three workflows described above (Figure 1).

Consistent with our expectations, different SL prediction approaches led to a diverse set of predicted SLIs. As shown in Figure S2, each workflow identified more than 1,000 synthetic lethal/synthetic dosage lethal partner genes, except for CGI, which identified only 67 synthetic lethal partner genes. Predicted SLI gene sets largely did not overlap; however, functional enrichment analysis showed shared pathway involvement in the interactions identified (Figure 3A; Table S2). In particular, we found significant KEGG pathway enrichment in synthetic lethal partner genes for the cell cycle, RNA metabolism, splicing machinery, chromatin organization, and transcriptional regulatory pathways (FDR < 0.05). Interestingly, several of these results are broadly related to genomic stability maintenance, and as such, confirm previously published reports from (Ku et al., 2020) and others that genes involved in SLIs tend to belong to related pathways. Gene ontology biological processes (GOBPs) enriched by synthetic lethal partner genes vary, but a clustering analysis based on hierarchical structure and semantic similarity of GOBPs showed the synthetic lethal partner genes identified via the different approaches were associated with similar biological processes (Figure 3B; Table S3). A clustering analysis summarized 350 GOBPs that were initially identified via four approaches into 25 representative GOBP groups. The 25 GOBP groups are associated with synthetic lethal partner genes identified by at least two workflows, and their biological functionality is mirrored by the pathway enrichment results presented above, with enrichment in genes involved in the cell cycle, transcriptional regulation, chromatin organization, and the DNA damage response. In summary, we have demonstrated that pathway-based SL prediction is easily implemented in this framework and can quickly generate useful insights beyond the single-gene level.

**Figure 3.**
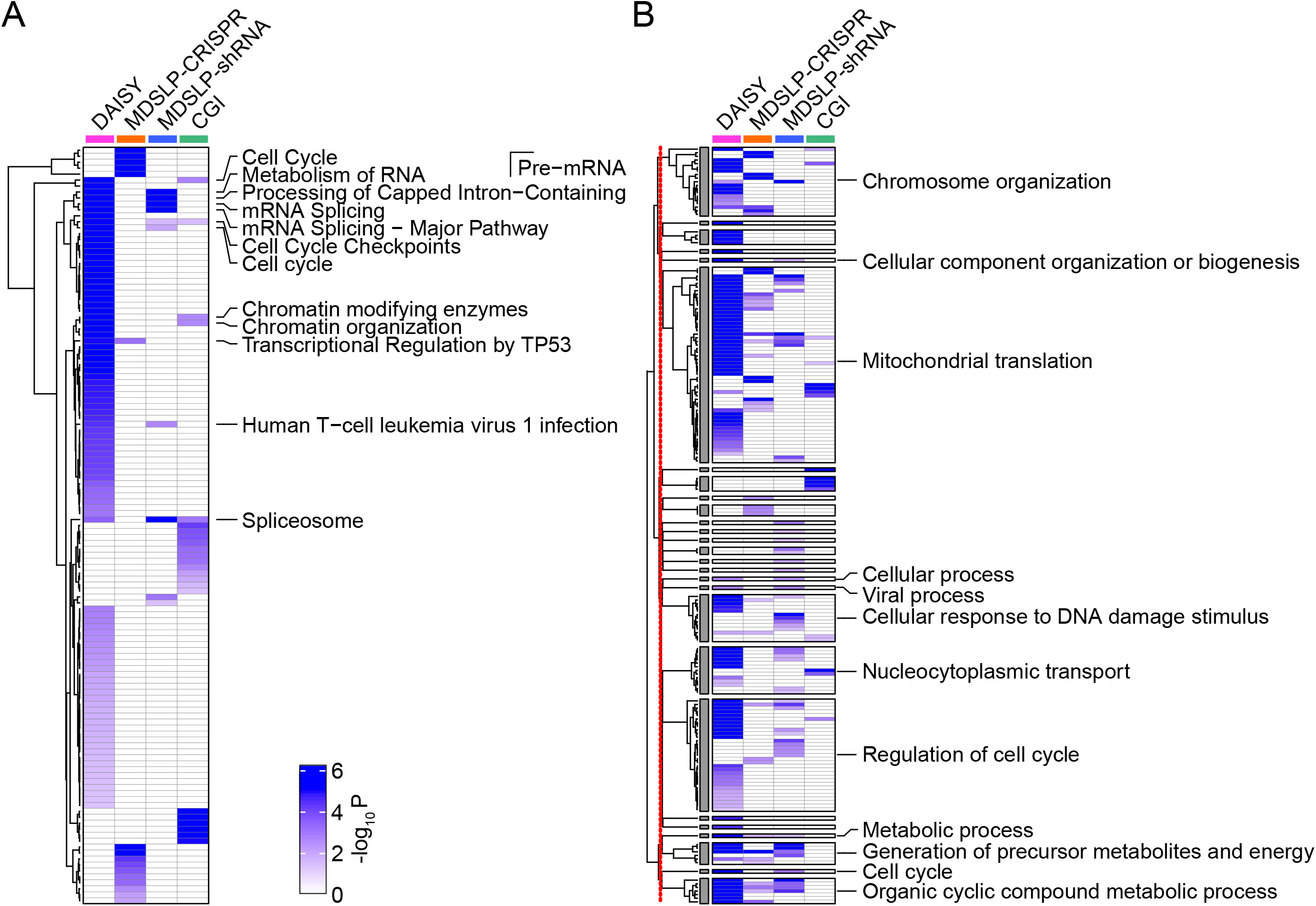
Pathway enrichment of predicted synthetic lethal partners of DNA damage repair (DDR) genes. Heatmaps depicting **A**. the KEGG or REACTOME pathway and **B**. Gene Ontology Biological Process (GOBP) enrichment for predictions made using four different approaches (columns). Increasing color intensity represents increasing statistical significance (*P* < 0.05, calculated by a hypergeometric test) for enrichment. Pathways or GOBPs were labelled if they were enriched by synthetic lethal partner genes identified by at least two prediction approaches. The redundant GOBPs were further reduced by REVIGO. The 398 GOBPs (Table S1) enriched by at least one approach were reduced to 172 GOBPs based on their semantic similarities and then summarized into 27 representative groups whose enrichment significance is represented in the heatmap. Clustering analysis was performed for GOBPs inside and outside of the 27 representative groups, separately. The red line down the left side of panel B indicates the separation between the clustering analyses. DAISY, data mining synthetic lethality identification pipeline; MDSLP-CRISPR, mutation-dependent synthetic lethality prediction pipeline with CRISPR; MDSLP-shRNA, mutation-dependent synthetic lethality prediction pipeline with shRNA; CGI, conserved synthetic lethal interactions from yeast screens

#### Tumor-specific SL analysis

Current evidence suggests that reproducible SLIs can be exquisitely context sensitive, e.g., being limited to a particular tissue type (Nijman and Friend, 2013; Ryan et al., 2018). We show that the MDSLP pipeline and our re-implementation of the DAISY algorithm can be applied to restricted subsets of the underlying data that represent samples or cell lines arising from the same cancer type. The rationale behind this type of analysis is that samples from the same cancer type could represent similar cellular origins, having a characteristic genetic interaction network that is tissue-type specific.

To illustrate this principle, we investigated the previously reported SLI between *ARID1A* and *ARID1B*. Functional loss of *ARID1B* is a specific vulnerability in *ARID1A*-mutated cancers, as it affects the composition of the SWI/SNF complex (Helming et al., 2014). In addition, protein levels of the core catalytic ATPase subunits such as SMARCA4, SMARCC2, and SMARCB1, were decreased with the deletion of *ARID1B* in an *ARID1A*-mutated ovarian cancer cell lines (Helming et al., 2014). We applied MDLSP and DAISY to finding synthetic lethal partners for *ARID1A*. Via the MDLSP pipeline, we found statistical evidence of differential dependency for *ARID1B* between *ARID1A*-mutated and wild-type cell lines in various cancer types, suggesting the potential for a SLI between the two genes across tissue types with the most reproducible SL potential in ovarian cancer (Figure 4A). Similar to our findings with the *BRCA*-related synthetic lethal partners, we also see differences in the strength of the relationship based on whether *ARID1B* was knocked down via shRNA or knocked out using the CRISPR-Cas9 system. The pan-cancer result is robust (shRNA/CRISPR), with more than 160 cell lines with *ARID1A* functional mutations. However, when constraining analysis to cell lines from a single cancer type, we have reduced power to detect differential effects due to limited sample size. Nevertheless, as there is strong and compelling evidence for this SLI, we still find support for the interaction occurring across multiple cancer types, even if the evidence comes from shRNA-derived or CRISPR-derived dependency datasets alone.

**Figure 4.**
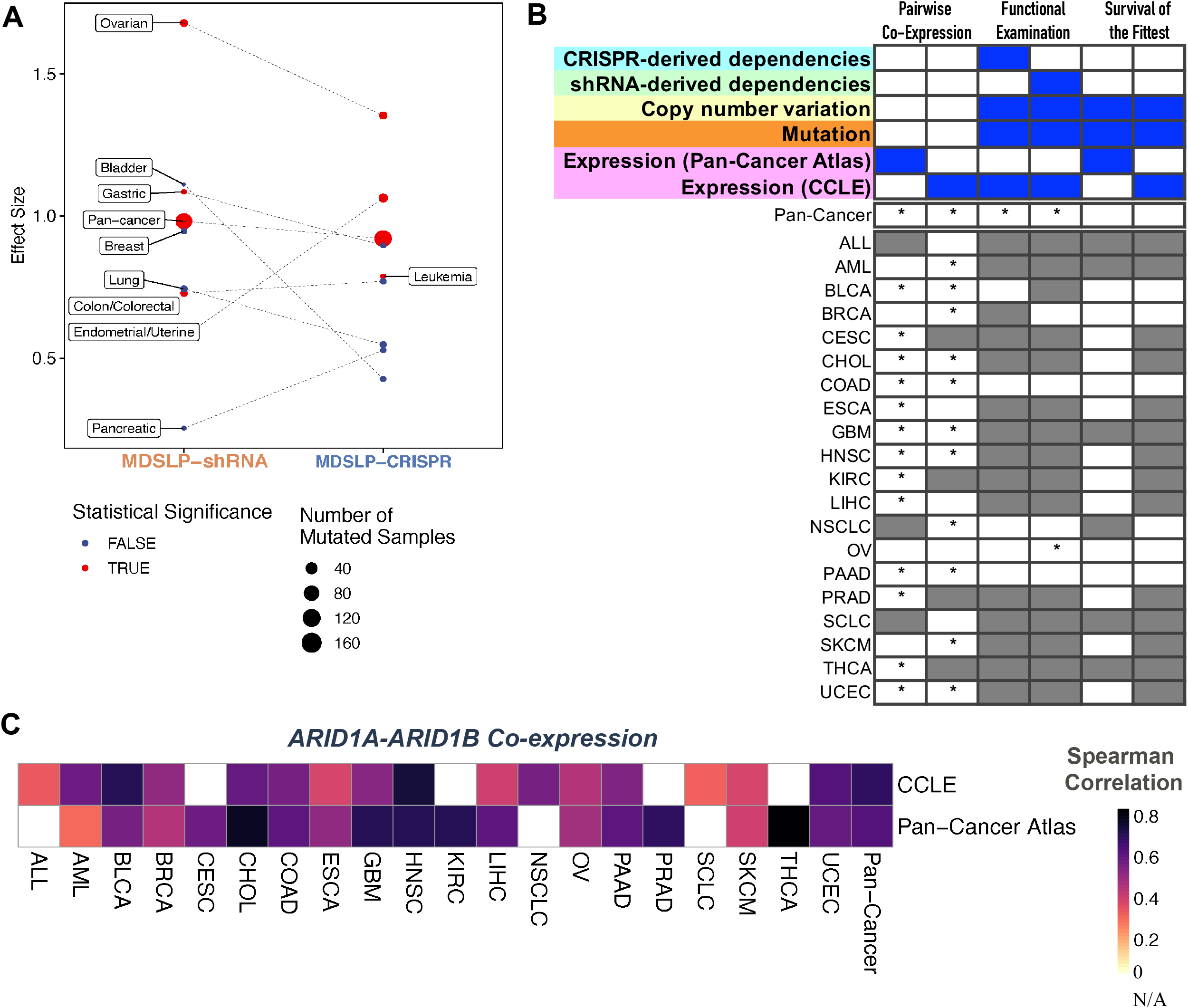
SL-Cloud enables cancer type-specific synthetic lethal inference for *ARID1A* and *ARID1B*. **A**. Evidence for synthetic lethality generated from the mutation-dependent synthetic lethality prediction (MDSLP) workflow as applied to cancer cell line dependency datasets when comparing the gene dependency scores (effects) for the shRNA and CRISPR datasets between the ARID1A-mutated group and wild-type group for different cancer types and across all cell lines (pan-cancer analysis). The threshold for statistical significance is FDR < 0.05. **B**. Statistical evidence for a synthetic lethal relationship between *ARID1A* and *ARID1B* from the data mining synthetic lethality identification pipeline (DAISY), with the results for each inference module being represented in the columns. Column heatmaps summarize the datasets used for each procedure. An asterisk (*) indicates that a test passed the FDR threshold (0.05); gray shading represents an invalid test or a lack of data availability. **C**. Heatmap visualization depicting the Spearman correlation between *ARID1A* and *ARID1B* for the annotated cancer type for a patient derived sample (Pan-Cancer Atlas) or cancer cell line (CCLE) (rows) across different cancer types (columns). Gray shading represents an invalid test.

DAISY does not predict an SLI between *ARID1A* and *ARID1B* when applied strictly, that is, when the requirement is set for statistically significant evidence across all three DAISY inference modules (Figure 4B). However, when considering each module individually, we see strong support for the interaction, with strong positive correlation (Spearman rho in the range [0.3, 0.77]) between these two genes across almost all cancer types considered (Figure 4B, C). Similar to the findings with MDSLP, we find evidence for the interaction between these two genes in ovarian cancer by using the functional examination module and shRNA-derived dependency dataset. This is unsurprising, as the underlying rationales for the DAISY and MDSLP inference strategies are quite similar and both approaches are applied to the same dataset. This further reinforces the tissue-specificity of the interaction, and is consistent with our expectations given the context in which that SLI was first described. We found no statistical support for the interaction via genomic SoF inference, which may be explained by the fact neither of those genes is inactivated by recurrent focal deletions that underpin that inference module.

Our findings suggest SL between *ARID1A* and *ARID1B* in some contexts but not in others. This illustrative example showcases the flexibility of the SL-Cloud framework in that it is relatively easy to compare the results of different SL prediction approaches, while varying algorithmic parameters, using the same or different data types, or to restrict analysis to a given tumor type for further elucidation of context specificities in SL.

## Discussion

The synthetic lethality concept presents a systematic framework with which to identify and nominate potential targets for cancer treatments. It is, therefore, critical to identify robust context-specific SLIs that provide insights into cancer vulnerabilities that can be nominated as targets for further therapeutic development. Although the SL concept offers a compelling rationale by which to inform drug target identification, systematically testing all potential SLIs in a given tissue or disease context is experimentally intractable. Therefore, it is necessary to develop reproducible computational inference and prioritization frameworks to nominate the most likely SLIs for experimental follow-up or to aid in functional screen design. We designed SL-Cloud to be a flexible framework to facilitate computational prediction of synthetic lethal genetic interactions to support investigations of potentially targetable cancer dependencies. We envisioned SL-Cloud as a tool with which computationally adept scientists could couple customized SL inferences pipelines and a seamless connection to large-scale datasets offered through cloud resources to facilitate customizable analysis, prediction, and nomination of novel SLIs.

SL-Cloud is designed to leverage existing cloud-based genomics infrastructure that takes advantage of the scalability of this technology and democratizes access to public large-scale cancer genomics datasets (Birger et al., 2017; Lau et al., 2017; Reynolds et al., 2017). In particular, SL-Cloud uses data resources and tools provided through the ISB-CGC, such as BigQuery, an SQL-based data warehousing solution that provides fast access through database-like queries. We have integrated publicly available omic and functional screening datasets for SL inference, and have demonstrated the key analytic steps needed to store analysis-ready processed data from data sources not currently hosted on the ISB-CGC. Users can analyze multiple large-scale datasets while circumventing the need to download and maintain a local copy of these petascale datasets or to perform extensive data management tasks. Overall, SL-Cloud offers an ensemble of methods and datasets that enables users to collate evidence for SLIs more easily, leveraging both the richness of existing publicly available datasets and facilitating the integration of smaller user-generated custom or private datasets into the same analysis framework.

There has been a concerted effort to develop novel SL prediction algorithms and to collect known and predicted SLIs in various databases (Deng et al., 2019; Guo et al., 2016; Han et al., 2019; Li et al., 2014; Wappett et al., 2021). In contrast to those approaches, we developed a framework for SLI discovery that is more flexible, that can incorporate state-of-the art algorithmic developments, and that can easily apply these methods to datasets generated with newer-generation screening technologies with increasing throughput. For example, our re-implementation of the DAISY pipeline is comprehensive in scope and expands on the initial report by using the most up-to-date versions of cloud-hosted datasets including patient-derived multi-omic cancer genomics datasets (TCGA) and cancer cell line characterization datasets (CCLE, shRNA, and CRISPR screens from DepMap) (Figure 1, Table S1), while giving the user the ability to adjust algorithmic settings and parameters.

Although the SL-Cloud framework is organized in a novel way that takes advantage of cloud-based data storage and computing, some of the limitations inherent in any one SL prediction strategy remain. Specifically, our ability to predict SLIs with high confidence is dependent on the size of the training dataset and on the underlying limitations of any one inference approach. For example, cancer type-specific analysis of the synthetic lethal potential of the *ARID1A/B* genes was in some cases limited by not having enough representative samples in a given cancer type with the target lesion in question.

In conclusion, we have presented here a community resource, SL-Cloud, that provides a practical framework to support investigations of context-specific SLIs. SL-Cloud is a unique implementation that brings together established SLI prediction workflows with the most up-to-date versions of large-scale multi-omics datasets to enable customizable SLI prediction. This organizational structure obviates the need for large local data downloads or extensive data management. So far, we have focused on axes such as prevalence of genomic alterations in human samples or interactions limited to specific pathways. However, the conceptual design of the framework is optimal for continued modular development and extensibility. We anticipate that this resource will enable investigators to integrate their own data with publicly available data to look for corroborating evidence for synthetic lethal genetic interactions with therapeutic potential and to explore such interactions in specific biological contexts.

## Acknowledgements

The research reported in this publication was supported by the National Cancer Institute of the National Institutes of Health under award number U01 CA217883 and P01CA077852. The results published here are fully or partially based upon data generated by the TCGA Research Network: (https://www.cancer.gov/tcga) and the Cancer Target Discovery and Development (CTD^2^) Network (https://ocg.cancer.gov/programs/ctd2/data-portal) established by the National Cancer Institute’s Office of Cancer Genomics. The content is solely the responsibility of the authors and does not necessarily represent the official views of the National Institutes of Health. The authors would like to thank the ISB-CGC Team for their support and Dr. William C. Hahn for facilitating access to and enabling the cloud-based redistribution of the Cancer Dependency Map data. We thank Dr. Arko Dasgupta and Russel Mosser from Fred Hutchinson Cancer Research Center for their constructive feedback, as well as many of our other colleagues and CTD^2^ collaborators for helpful discussions and critical manuscript feedback. The authors thank Keith A. Laycock, PhD, ELS, for scientific editing of the manuscript. We dedicate this paper to the memory of Dr. Daniela S. Gerhard. We are grateful for her constant encouragement, scientific suggestions and support for this project.

## Author Contributions

Conceptualization, B.T., G.Q., T.K., N.C. and I.S; Methodology, Investigation, Formal Analysis and Software, B.T. G.Q. T.K. and B.A.; Data Curation, B.T. and T.K.; Writing - Original Draft, B.T. G.Q, T.K and N.C.; Writing - Review & Editing, B.T. G.Q, T.K, B.A., C.J.K, N.C. and I.S.; Funding Acquisition, C.J.K, I.S.; Resources, B.A.; Supervision, N.C. and I.S.;

## Figure Legends

**Figure S1.**
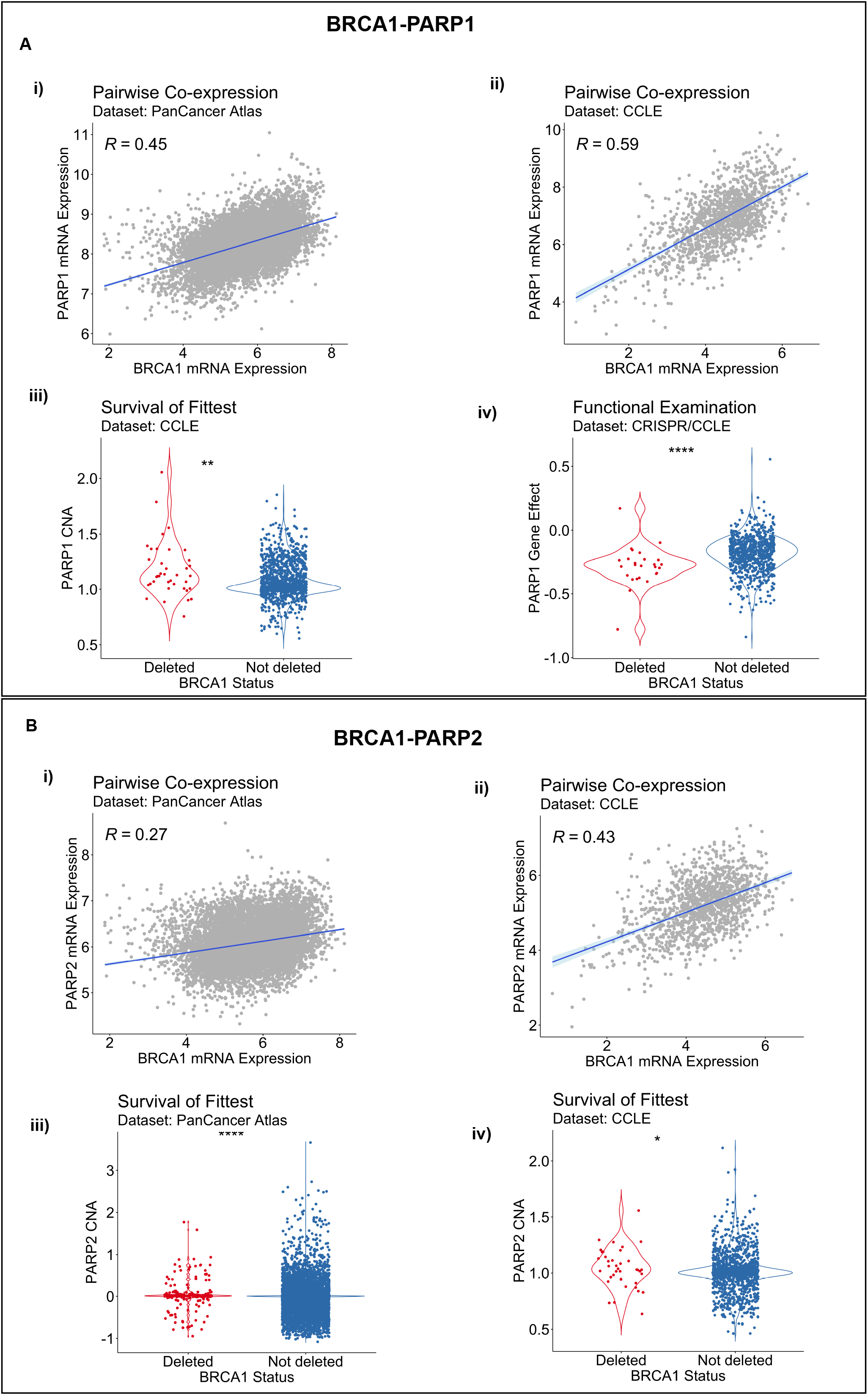
Evidence for BRCA1-PARP1/2 synthetic lethal relationship based on the data mining synthetic lethality identification pipeline (DAISY). **A**. Evidence for a *BRCA1-PARP1* synthetic lethal relationship by pairwise co-expression in i) Pan-Cancer Atlas and ii) CCLE datasets respectively; iii) by survival of the fittest in in CCLE data; and iv) by functional examination in DepMap CRISPR. **B**. Evidence for a *BRCA1-PARP2* SL relationship by pairwise co-expression in i) Pan-Cancer Atlas and ii) CCLE datasets and by survival of the fittest in iii) Pan-Cancer Atlas and iv) CCLE datasets. *R*, Spearman correlation coefficient; p, P value by the Wilcoxon rank-sum test; * *P* < 0.05; ** *P* <0.01; **** *P* < 0.0001.

**Figure S1.**
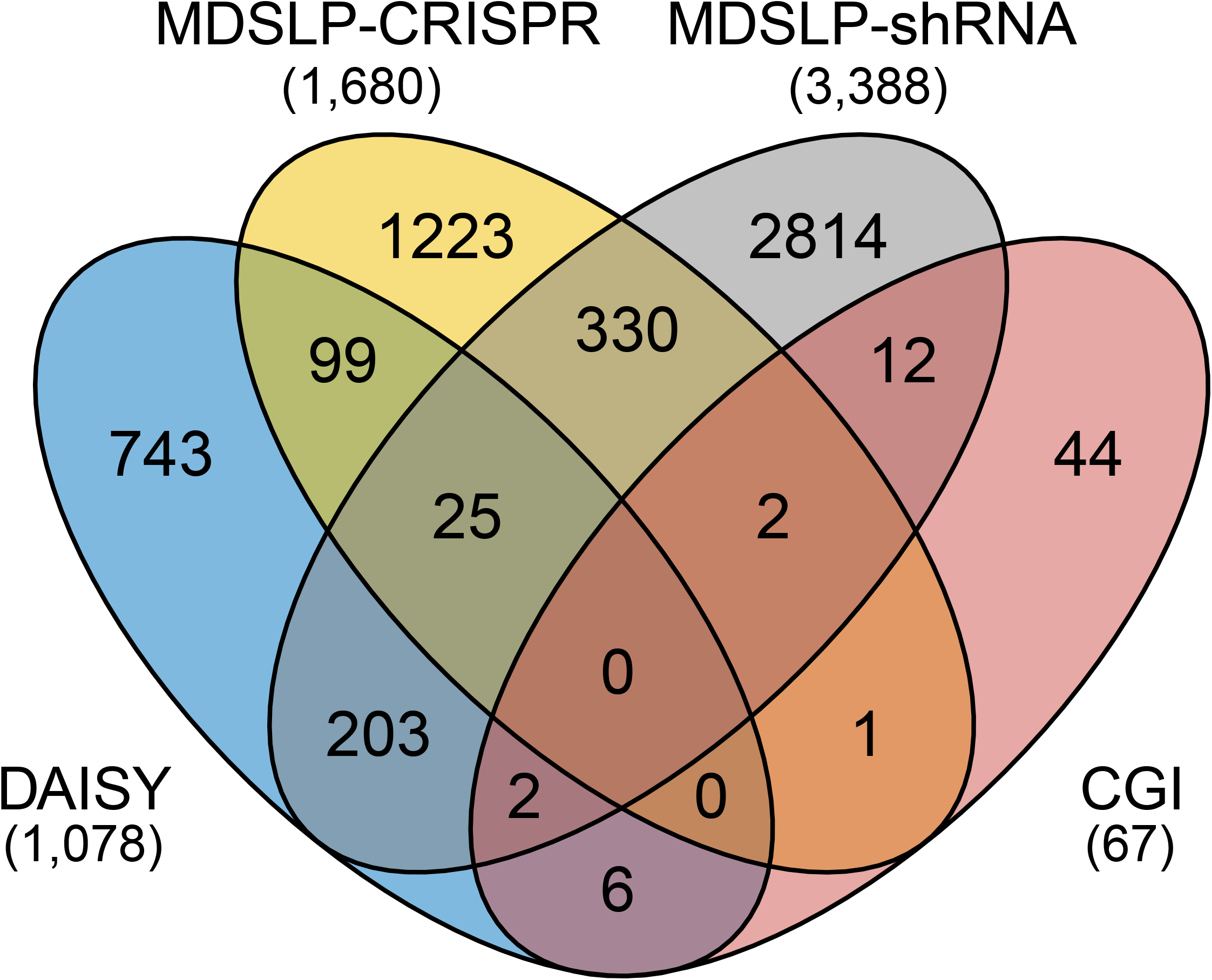
Relatedness of synthetic lethal partners of DDR genes as predicted by different approaches. Each set represents the number of synthetic lethal partner genes identified by different approaches. The numbers in parentheses are the numbers of partner genes predicted by each approach. MDSLP, mutation-dependent synthetic lethality prediction; CGI, conserved genetic interactions.

## Table Legends

**Table S1**. Publicly available cancer genomic and molecular profiling datasets relevant for SL inference

**Table S2**. Pathways enriched by synthetic lethal partners of DDR genes

**Table S3**. Gene ontology biological processes enriched by synthetic lethal partners of DDR genes

