## Supplemental Table 1 for "SL-Cloud: A Computational Resource to Support Synthetic Lethal Interaction Discovery"

Table S1. Publicly available cancer genomic and molecular profilling datasets relevant for SL inference

| Data | Data Resource | Data Type | Data/Table | Google Bigquery Table ID | Approach | Reference | Link (Original Data Resource Address) |
|---|---|---|---|---|---|---|---|
| TCGA omics data | TCGA (The Cancer Genome Atlas) | Copy number variation | all_data_by_genes_whitelisted.tsv | isb-cgc-bq.pancancer_atlas.Filtered_all_CNVR_data_by_gene | DAISY | Hutter and Zenklusen, 2018 | https://www.synapse.org/#!Synapse:syn5049514 |
| | | Gene expression | EBPlusPlusAdjustPANCAN_IlluminaHiSeq_RNASeqV2.geneExp.tsv | isb-cgc-bq.pancancer_atlas.Filtered_EBpp_AdjustPANCAN_IlluminaHiSeq_RNASeqV2_genExp | DAISY | | https://api.gdc.cancer.gov/data/3586c0da-64d0-4b74-a449-5ff4d9136611 |
| | | Mutation | Pancan.merged.v0.2.5.filtered.maf.gz | isb-cgc-bq.pancancer_atlas.Filtered_MC3_MAF_V5_one_per_tumor_sample | DAISY | | https://api.gdc.cancer.gov/data/c946eefc-20a0-4277-a6da-fe42ed4d793a |
| CCLE omics data | CCLE (Cancer Cell Line Encyclopedia) | Copy number variation | CCLE_gene_cn.csv | syntheticlethality.DepMap_public_20Q3.CCLE_gene_cn | DAISY | Ghandi et al. 2019 | https://ndownloader.figshare.com/files/24613352 |
| | | Gene expression | CCLE_expression.csv | syntheticlethality.DepMap_public_20Q3.CCLE_gene_expression | DAISY | | https://ndownloader.figshare.com/files/24613325 |
| | | Mutation | CCLE_mutations.csv | syntheticlethality.DepMap_public_20Q3.CCLE_mutation | DAISY / MDSLP | | https://ndownloader.figshare.com/files/24613355 |
| | | Sample information | sample_info.csv | syntheticlethality.DepMap_public_20Q3.sample_info_Depmap_withTCGA_labels | DAISY / MDSLP | | https://ndownloader.figshare.com/files/24613394 |
| CRISPR based gene effects | DepMap (The Cancer Dependency Map) | Gene dependency score | D2_Achilles_gene_effect.csv | syntheticlethality:DepMap_public_20Q3.Achilles_gene_effect | DAISY / MDSLP | Meyet et al. 2017, Dempster et al. 2019, Ghandi et al. 2019, Broad 2020 | https://ndownloader.figshare.com/files/24613292/ |
| shRNA based gene effects | Achilles DRIVE Marcotte et. al. 2016 | Gene dependency score | D2_combined_gene_dep_scores.csv | syntheticlethality.DEMETER2_v6.D2_combined_gene_dep_score | DAISY / MDSLP | McFarland et al. 2018 | https://ndownloader.figshare.com/files/13515395 |
| Genetic interactions in Yeast | TheCellMap | Genetic interaction scores | Raw genetic interaction datasets: Pair-wise interaction format.zip | syntheticlethality.CellMap.CellMap | CGI | Costanzo et al. 2016 | https://thecellmap.org/costanzo2016/ |
