## Supplemental Table 2 for "SL-Cloud: A Computational Resource to Support Synthetic Lethal Interaction Discovery"

**Table S2.** Pathways enriched by synthetic lethal partners of DDR genes

| Pathway id | Pathway | DAISY | MDSLP-CR | MDSLP-shF | CGI |
|---|---|---|---|---|---|
| KEGG:00310 | Lysine degradation | 0.046 | | | |
| KEGG:00970 | Aminoacyl-tRNA biosynthesis | | 0.001 | | |
| KEGG:01522 | Endocrine resistance | 0.014 | | | |
| KEGG:03010 | Ribosome | | 0.000 | | |
| KEGG:03013 | RNA transport | 0.001 | | | |
| KEGG:03015 | mRNA surveillance pathway | 0.000 | | | |
| KEGG:03022 | Basal transcription factors | 0.038 | | | |
| KEGG:03040 | Spliceosome | 0.000 | | 0.000 | 0.001 |
| KEGG:04010 | MAPK signaling pathway | | | | 0.045 |
| KEGG:04068 | FoxO signaling pathway | 0.020 | | | |
| KEGG:04110 | Cell cycle | 0.000 | | 0.012 | |
| KEGG:04114 | Oocyte meiosis | 0.005 | | | |
| KEGG:04141 | Protein processing in endoplasmic reticulum | | | | 0.002 |
| KEGG:04144 | Endocytosis | | | | 0.000 |
| KEGG:04213 | Longevity regulating pathway - multiple species | | | | 0.000 |
| KEGG:04218 | Cellular senescence | 0.000 | | | |
| KEGG:04612 | Antigen processing and presentation | | | | 0.000 |
| KEGG:04714 | Thermogenesis | | 0.003 | | |
| KEGG:04914 | Progesterone-mediated oocyte maturation | 0.000 | | | |
| KEGG:04915 | Estrogen signaling pathway | | | | 0.001 |
| KEGG:04932 | Non-alcoholic fatty liver disease | | 0.009 | | |
| KEGG:05020 | Prion disease | | | | 0.030 |
| KEGG:05134 | Legionellosis | | | | 0.000 |
| KEGG:05145 | Toxoplasmosis | | | | 0.000 |
| KEGG:05161 | Hepatitis B | 0.000 | | | |
| KEGG:05162 | Measles | | | | 0.001 |
| KEGG:05166 | Human T-cell leukemia virus 1 infection | 0.000 | | 0.002 | |
| KEGG:05167 | Kaposi sarcoma-associated herpesvirus infection | 0.024 | | | |
| KEGG:05169 | Epstein-Barr virus infection | 0.002 | | | |
| KEGG:05170 | Human immunodeficiency virus 1 infection | 0.013 | | | |
| KEGG:05203 | Viral carcinogenesis | 0.018 | | | |
| KEGG:05211 | Renal cell carcinoma | 0.005 | | | |
| KEGG:05215 | Prostate cancer | 0.000 | | | |
| KEGG:05220 | Chronic myeloid leukemia | 0.004 | | | |
| KEGG:05222 | Small cell lung cancer | 0.035 | | | |
| KEGG:05223 | Non-small cell lung cancer | 0.040 | | | |
| KEGG:05417 | Lipid and atherosclerosis | | | | 0.008 |
| REAC:R-HSA | E2F-enabled inhibition of pre-replication complex f | 0.048 | | | |
| REAC:R-HSA | E2F mediated regulation of DNA replication | 0.002 | | | |
| REAC:R-HSA | Transcription of E2F targets under negative contro | 0.046 | | | |
| REAC:R-HSA | Transcription of E2F targets under negative contro | 0.012 | | | |
| REAC:R-HSA | Amplification of signal from the kinetochores | 0.000 | | | |
| REAC:R-HSA | Amplification of signal from unattached kinetocho | 0.000 | | | |
| REAC:R-HSA | The citric acid (TCA) cycle and respiratory electron transport | | 0.000 | | |

| | | | | | |
|---|---|---|---|---|---|
| REAC:R-HSA | G0 and Early G1 | 0.000 | | | |
| REAC:R-HSA | Polo-like kinase mediated events | 0.000 | | | |
| REAC:R-HSA | Transport of Mature mRNA derived from an Intron- | 0.000 | | | |
| REAC:R-HSA | Respiratory electron transport, ATP synthesis by chemiosmotic | | 0.001 | | |
| REAC:R-HSA | Cell Cycle | 0.000 | | | 0.002 |
| REAC:R-HSA | APC/C-mediated degradation of cell cycle proteins | 0.012 | | | |
| REAC:R-HSA | Activation of ATR in response to replication stress | 0.001 | | | |
| REAC:R-HSA | Unwinding of DNA | 0.001 | | | |
| REAC:R-HSA | RHO GTPase Effectors | 0.003 | | | |
| REAC:R-HSA | Generic Transcription Pathway | 0.000 | | | |
| REAC:R-HSA | Separation of Sister Chromatids | 0.000 | | | |
| REAC:R-HSA | Resolution of Sister Chromatid Cohesion | 0.000 | | | |
| REAC:R-HSA | Mitotic Metaphase and Anaphase | 0.000 | | | |
| REAC:R-HSA | SUMOylation | 0.002 | | | |
| REAC:R-HSA | SUMO E3 ligases SUMOylate target proteins | 0.003 | | | |
| REAC:R-HSA | PKMTs methylate histone lysines | 0.050 | | | |
| REAC:R-HSA | HDMs demethylate histones | | | | 0.000 |
| REAC:R-HSA | Chromatin modifying enzymes | 0.000 | | | 0.003 |
| REAC:R-HSA | Regulation of HSF1-mediated heat shock response | | | | 0.000 |
| REAC:R-HSA | HSP90 chaperone cycle for steroid hormone receptors (SHR | | | | 0.001 |
| REAC:R-HSA | Cellular response to heat stress | | | | 0.000 |
| REAC:R-HSA | Attenuation phase | | | | 0.000 |
| REAC:R-HSA | HSF1-dependent transactivation | | | | 0.000 |
| REAC:R-HSA | Transcriptional Regulation by TP53 | 0.000 | 0.001 | | |
| REAC:R-HSA | tRNA Aminoacylation | | 0.008 | | |
| REAC:R-HSA | Mitochondrial tRNA aminoacylation | | 0.000 | | |
| REAC:R-HSA | Mitotic G2-G2/M phases | 0.000 | | | |
| REAC:R-HSA | Regulation of mitotic cell cycle | 0.012 | | | |
| REAC:R-HSA | Mitotic G1 phase and G1/S transition | 0.000 | | | |
| REAC:R-HSA | Chromatin organization | 0.000 | | | 0.003 |
| REAC:R-HSA | Mitochondrial translation initiation | | 0.000 | | |
| REAC:R-HSA | Mitochondrial translation | | 0.000 | | |
| REAC:R-HSA | Mitochondrial translation elongation | | 0.000 | | |
| REAC:R-HSA | Mitochondrial translation termination | | 0.000 | | |
| REAC:R-HSA | TP53 Regulates Metabolic Genes | | 0.000 | | |
| REAC:R-HSA | Regulation of TP53 Activity | 0.000 | | | |
| REAC:R-HSA | RHO GTPases Activate Formins | 0.000 | | | |
| REAC:R-HSA | HDR through Single Strand Annealing (SSA | | | 0.045 | |
| REAC:R-HSA | Respiratory electron transport | | 0.000 | | |
| REAC:R-HSA | TP53 Regulates Transcription of Cell Cycle Genes | 0.000 | | | |
| REAC:R-HSA | Regulation of TP53 Activity through Acetylation | 0.029 | | | |
| REAC:R-HSA | PTEN Regulation | 0.006 | | | |
| REAC:R-HSA | CDC6 association with the ORC:origin complex | 0.000 | | | |
| REAC:R-HSA | Assembly of the pre-replicative complex | 0.000 | | | |
| REAC:R-HSA | Mitotic Prometaphase | 0.000 | | | |

| | | | | | |
|---|---|---|---|---|---|
| REAC:R-HSA | Mitotic Anaphase | 0.000 | | | |
| REAC:R-HSA | Mitotic Telophase/Cytokinesis | 0.002 | | | |
| REAC:R-HSA | M Phase | 0.000 | | | |
| REAC:R-HSA | Orc1 removal from chromatin | 0.001 | | | |
| REAC:R-HSA | Activation of the pre-replicative complex | 0.000 | | | |
| REAC:R-HSA | DNA Replication Pre-Initiation | 0.000 | | | |
| REAC:R-HSA | Switching of origins to a post-replicative state | 0.000 | | | |
| REAC:R-HSA | DNA strand elongation | 0.000 | | | |
| REAC:R-HSA | Cyclin E associated events during G1/S transition | 0.021 | | | |
| REAC:R-HSA | G1/S-Specific Transcription | 0.000 | | | |
| REAC:R-HSA | G1/S Transition | 0.000 | | | |
| REAC:R-HSA | Synthesis of DNA | 0.000 | | | |
| REAC:R-HSA | S Phase | 0.000 | | | |
| REAC:R-HSA | G2/M Transition | 0.000 | | | |
| REAC:R-HSA | Cell Cycle, Mitotic | 0.000 | | | |
| REAC:R-HSA | DNA Replication | 0.000 | | | |
| REAC:R-HSA | G2/M Checkpoints | | | | 0.014 |
| REAC:R-HSA | Mitotic Spindle Checkpoint | 0.000 | | | |
| REAC:R-HSA | Cell Cycle Checkpoints | 0.000 | | 0.034 | 0.037 |
| REAC:R-HSA | Cyclin A:Cdk2-associated events at S phase entry | 0.029 | | | |
| REAC:R-HSA | mRNA Splicing - Major Pathway | 0.000 | | 0.000 | |
| REAC:R-HSA | mRNA Splicing - Minor Pathway | | | 0.001 | |
| REAC:R-HSA | mRNA Splicing | 0.000 | | 0.000 | |
| REAC:R-HSA | mRNA 3'-end processing | 0.000 | | | |
| REAC:R-HSA | Transport of Mature Transcript to Cytoplasm | 0.000 | | | |
| REAC:R-HSA | Processing of Capped Intron-Containing Pre-mRN | 0.000 | | 0.000 | |
| REAC:R-HSA | Translation | | 0.000 | | |
| REAC:R-HSA | RNA Polymerase II Transcription Termination | 0.000 | | | |
| REAC:R-HSA | RNA Polymerase II Transcription | 0.000 | | | |
| REAC:R-HSA | Chromosome Maintenance | 0.007 | | | |
| REAC:R-HSA | Gene expression (Transcription | 0.000 | | | |
| REAC:R-HSA | ESR-mediated signaling | 0.037 | | | |
| REAC:R-HSA | Metabolism of RNA | 0.000 | | 0.000 | |
| REAC:R-HSA | EML4 and NUDC in mitotic spindle formation | 0.000 | | | |
| REAC:R-HSA | Aberrant regulation of mitotic G1/S transition in car | 0.020 | | | |
| REAC:R-HSA | Defective binding of RB1 mutants to E2F1,(E2F2, | 0.020 | | | |
| REAC:R-HSA | Diseases of mitotic cell cycle | 0.039 | | | |
| REAC:R-HSA | Aberrant regulation of mitotic cell cycle due to RB1 | 0.024 | | | |
