## Supplemental Table 3 for "SL-Cloud: A Computational Resource to Support Synthetic Lethal Interaction Discovery"

**Table S3.** Gene ontology biological processes enriched by synthetic lethal partners of DDR genes

| GO id | GO term | Representative GO term | DAISY | MDSLP-CR | MDSLP-sh | CGI |
|---|---|---|---|---|---|---|
| GO:0007568 | aging | aging | | | 8.089E-04 | |
| GO:0007569 | cell aging | aging | | | 3.446E-02 | |
| GO:0008152 | metabolic process | metabolic process | 3.866E-24 | 1.630E-02 | 2.368E-02 | |
| GO:0009987 | cellular process | cellular process | 2.134E-03 | | 3.817E-03 | |
| GO:0016032 | viral process | viral process | 2.120E-03 | | 2.081E-03 | |
| GO:0023052 | signaling | signaling | | | 3.839E-02 | |
| GO:0032543 | mitochondrial translation | mitochondrial translation | | 4.748E-53 | | |
| GO:0043038 | amino acid activation | mitochondrial translation | | 3.195E-02 | | |
| GO:0044265 | cellular macromolecule catabolic p | mitochondrial translation | 5.723E-08 | | | 4.246E-02 |
| GO:0044260 | cellular macromolecule metabolic | mitochondrial translation | 2.188E-33 | | | |
| GO:0018205 | peptidyl-lysine modification | mitochondrial translation | 8.003E-11 | | | |
| GO:0140053 | mitochondrial gene expression | mitochondrial translation | | 3.182E-53 | | |
| GO:0006482 | protein demethylation | mitochondrial translation | | | | 5.192E-10 |
| GO:0008214 | protein dealkylation | mitochondrial translation | | | | 5.192E-10 |
| GO:0006351 | transcription DNA-templated | mitochondrial translation | 2.190E-18 | | | |
| GO:0006270 | DNA replication initiation | mitochondrial translation | 7.354E-08 | | | |
| GO:0006473 | protein acetylation | mitochondrial translation | 3.320E-06 | | | |
| GO:0043543 | protein acylation | mitochondrial translation | 4.356E-05 | | | |
| GO:0000959 | mitochondrial RNA metabolic proc | mitochondrial translation | | 1.402E-06 | | |
| GO:0043412 | macromolecule modification | mitochondrial translation | 2.126E-12 | | | |
| GO:0009059 | macromolecule biosynthetic proces | mitochondrial translation | 1.425E-24 | 1.235E-02 | | |
| GO:0010467 | gene expression | mitochondrial translation | 7.543E-25 | 2.366E-03 | | |
| GO:0008213 | protein alkylation | mitochondrial translation | 1.741E-03 | | | |
| GO:0006139 | nucleobase-containing compound | mitochondrial translation | 1.221E-49 | | 1.859E-06 | |
| GO:0044267 | cellular protein metabolic process | mitochondrial translation | 1.234E-09 | 1.166E-02 | | |
| GO:0070647 | protein modification by small prote | mitochondrial translation | 3.404E-08 | | | |
| GO:0043603 | cellular amide metabolic process | mitochondrial translation | | 4.496E-12 | | |
| GO:0043414 | macromolecule methylation | mitochondrial translation | 2.796E-05 | | | |
| GO:0090305 | nucleic acid phosphodiester bond | mitochondrial translation | 4.745E-04 | | | |
| GO:0019538 | protein metabolic process | mitochondrial translation | 5.896E-05 | 2.413E-02 | | |
| GO:0006260 | DNA replication | mitochondrial translation | 4.975E-09 | | | |
| GO:0032446 | protein modification by small prote | mitochondrial translation | 1.439E-04 | | | |
| GO:0006310 | DNA recombination | mitochondrial translation | 5.382E-05 | | | |
| GO:0006397 | mRNA processing | mitochondrial translation | 1.120E-13 | 1.861E-02 | 1.262E-04 | |
| GO:0044271 | cellular nitrogen compound biosynt | mitochondrial translation | 3.883E-23 | 1.141E-04 | | |
| GO:0018193 | peptidyl-amino acid modification | mitochondrial translation | 5.678E-06 | | | |
| GO:0016071 | mRNA metabolic process | mitochondrial translation | 6.675E-15 | | 6.415E-04 | 2.006E-02 |
| GO:0031123 | RNA 3'-end processing | mitochondrial translation | 1.214E-05 | | | |
| GO:0016070 | RNA metabolic process | mitochondrial translation | 2.372E-34 | | 2.890E-03 | |
| GO:1901362 | organic cyclic compound biosynthe | mitochondrial translation | 3.834E-20 | | | |
| GO:0019438 | aromatic compound biosynthetic pr | mitochondrial translation | 2.086E-21 | | | |
| GO:0018130 | heterocycle biosynthetic process | mitochondrial translation | 1.317E-21 | | | |
| GO:0031145 | anaphase-promoting complex-depe | mitochondrial translation | 3.554E-02 | | | |
| GO:0018022 | peptidyl-lysine methylation | mitochondrial translation | 4.015E-04 | | | |
| GO:1901576 | organic substance biosynthetic pro | mitochondrial translation | 1.515E-20 | 4.411E-04 | | |
| GO:0006259 | DNA metabolic process | mitochondrial translation | 1.484E-25 | | 1.081E-03 | |
| GO:0034645 | cellular macromolecule biosyntheti | mitochondrial translation | 1.546E-24 | 1.053E-02 | | |

| | | | | | | |
|---|---|---|---|---|---|---|
| GO:1901566 | organonitrogen compound biosynth | mitochondrial translation | | 3.311E-11 | | |
| GO:0008380 | RNA splicing | mitochondrial translation | 5.421E-12 | | 3.714E-05 | |
| GO:0006396 | RNA processing | mitochondrial translation | 3.758E-18 | 4.121E-05 | 1.283E-06 | |
| GO:1901361 | organic cyclic compound catabolic | mitochondrial translation | | | | 1.204E-04 |
| GO:0006468 | protein phosphorylation | mitochondrial translation | | | 2.101E-02 | |
| GO:0006796 | phosphate-containing compound m | mitochondrial translation | | | 1.402E-04 | |
| GO:0090304 | nucleic acid metabolic process | mitochondrial translation | 8.006E-51 | | 4.967E-05 | |
| GO:0032774 | RNA biosynthetic process | mitochondrial translation | 2.874E-19 | | | |
| GO:0006399 | tRNA metabolic process | mitochondrial translation | | 6.362E-03 | | |
| GO:0034654 | nucleobase-containing compound | mitochondrial translation | 3.670E-21 | | | |
| GO:0034655 | nucleobase-containing compound | mitochondrial translation | 1.534E-03 | | | 1.902E-05 |
| GO:0009057 | macromolecule catabolic process | mitochondrial translation | 3.597E-07 | | | |
| GO:0006261 | DNA-dependent DNA replication | mitochondrial translation | 3.324E-06 | | | |
| GO:0050896 | response to stimulus | response to stimulus | | | 1.495E-03 | |
| GO:0051726 | regulation of cell cycle | regulation of cell cycle | 4.257E-15 | | 7.381E-03 | |
| GO:0031647 | regulation of protein stability | regulation of cell cycle | 3.325E-04 | | | |
| GO:1901796 | regulation of signal transduction by | regulation of cell cycle | 9.520E-05 | | | |
| GO:0051983 | regulation of chromosome segrega | regulation of cell cycle | 2.337E-07 | | | |
| GO:0033044 | regulation of chromosome organiza | regulation of cell cycle | 4.754E-10 | | | |
| GO:0010564 | regulation of cell cycle process | regulation of cell cycle | 3.996E-15 | | | |
| GO:0051173 | positive regulation of nitrogen com | regulation of cell cycle | 1.827E-12 | 4.640E-03 | 4.729E-05 | |
| GO:0062125 | regulation of mitochondrial gene ex | regulation of cell cycle | | 3.304E-03 | | |
| GO:0051128 | regulation of cellular component or | regulation of cell cycle | 1.036E-02 | | | |
| GO:0048518 | positive regulation of biological pro | regulation of cell cycle | | | 6.903E-05 | |
| GO:0040029 | regulation of gene expression epig | regulation of cell cycle | 1.370E-02 | | | |
| GO:0065008 | regulation of biological quality | regulation of cell cycle | | | 3.398E-03 | |
| GO:0043484 | regulation of RNA splicing | regulation of cell cycle | 1.689E-02 | | | |
| GO:1903311 | regulation of mRNA metabolic proc | regulation of cell cycle | 2.233E-09 | | | 1.775E-03 |
| GO:0051052 | regulation of DNA metabolic proce | regulation of cell cycle | 3.169E-06 | | 4.357E-02 | |
| GO:0031329 | regulation of cellular catabolic proc | regulation of cell cycle | 6.472E-04 | | | |
| GO:0009894 | regulation of catabolic process | regulation of cell cycle | 4.165E-05 | | | |
| GO:0019222 | regulation of metabolic process | regulation of cell cycle | 2.030E-14 | | | |
| GO:0006357 | regulation of transcription by RNA | regulation of cell cycle | 3.881E-10 | | | |
| GO:0034248 | regulation of cellular amide metabo | regulation of cell cycle | 1.779E-02 | | | |
| GO:0045934 | negative regulation of nucleobase- | regulation of cell cycle | 2.886E-08 | | 4.872E-03 | |
| GO:0010608 | posttranscriptional regulation of ge | regulation of cell cycle | 4.879E-02 | | | |
| GO:0032268 | regulation of cellular protein metab | regulation of cell cycle | 9.323E-04 | | | |
| GO:0051246 | regulation of protein metabolic proc | regulation of cell cycle | 3.883E-03 | | | |
| GO:0070129 | regulation of mitochondrial translat | regulation of cell cycle | | 4.379E-03 | | |
| GO:1901216 | positive regulation of neuron death | regulation of cell cycle | | | 2.401E-03 | |
| GO:1900182 | positive regulation of protein localiz | regulation of cell cycle | 1.550E-02 | | | |
| GO:0019219 | regulation of nucleobase-containin | regulation of cell cycle | 4.025E-22 | | 2.583E-02 | |
| GO:1902531 | regulation of intracellular signal tra | regulation of cell cycle | | | 1.525E-03 | |
| GO:1902275 | regulation of chromatin organizatio | regulation of cell cycle | 8.865E-04 | | | |
| GO:0050794 | regulation of cellular process | regulation of cell cycle | | | 2.642E-03 | |
| GO:0065007 | biological regulation | biological regulation | | | 1.362E-02 | |
| GO:0051276 | chromosome organization | chromosome organization | 5.450E-48 | | | 1.211E-02 |
| GO:0006415 | translational termination | chromosome organization | | 6.366E-37 | | |

| | | | | | | |
|---|---|---|---|---|---|---|
| GO:0006414 | translational elongation | chromosome organization | | 2.852E-33 | | |
| GO:0051383 | kinetochore organization | chromosome organization | 3.781E-03 | | | |
| GO:0022411 | cellular component disassembly | chromosome organization | | 1.388E-12 | | |
| GO:0022613 | ribonucleoprotein complex biogene | chromosome organization | 4.597E-03 | | 9.430E-06 | |
| GO:0000723 | telomere maintenance | chromosome organization | 9.138E-04 | | | |
| GO:0032200 | telomere organization | chromosome organization | 1.389E-03 | | | |
| GO:0006323 | DNA packaging | chromosome organization | 3.068E-06 | | | |
| GO:0043933 | protein-containing complex subunit | chromosome organization | 1.225E-04 | 9.748E-05 | | |
| GO:0000280 | nuclear division | chromosome organization | 1.424E-15 | | | |
| GO:0043604 | amide biosynthetic process | chromosome organization | | 1.070E-16 | | |
| GO:0048285 | organelle fission | chromosome organization | 3.316E-15 | | | |
| GO:0007005 | mitochondrion organization | chromosome organization | | 1.503E-05 | | |
| GO:0071824 | protein-DNA complex subunit orga | chromosome organization | 1.358E-07 | | | |
| GO:0065004 | protein-DNA complex assembly | chromosome organization | 1.060E-05 | | | |
| GO:0033108 | mitochondrial respiratory chain cor | chromosome organization | | 1.878E-03 | | |
| GO:0006996 | organelle organization | chromosome organization | 3.532E-22 | | | 2.621E-04 |
| GO:0006325 | chromatin organization | chromosome organization | 2.150E-26 | | | |
| GO:0042026 | protein refolding | protein refolding | | | | 2.938E-07 |
| GO:0051085 | chaperone cofactor-dependent pro | protein refolding | | | | 2.096E-06 |
| GO:0006458 | de novo' protein folding | protein refolding | | | | 1.066E-05 |
| GO:0061077 | chaperone-mediated protein foldin | protein refolding | | | | 1.772E-04 |
| GO:0008219 | cell death | cell death | | | 2.556E-02 | |
| GO:0006913 | nucleocytoplasmic transport | nucleocytoplasmic transpo | 5.524E-11 | | 4.739E-04 | |
| GO:0007041 | lysosomal transport | nucleocytoplasmic transport | | | | 2.489E-04 |
| GO:0031503 | protein-containing complex localiza | nucleocytoplasmic transpo | 8.230E-04 | | | |
| GO:0051028 | mRNA transport | nucleocytoplasmic transpo | 3.252E-10 | | | |
| GO:0007034 | vacuolar transport | nucleocytoplasmic transport | | | | 2.990E-07 |
| GO:0033036 | macromolecule localization | nucleocytoplasmic transport | | | 1.946E-02 | |
| GO:0051641 | cellular localization | nucleocytoplasmic transport | | | 4.305E-02 | |
| GO:0015931 | nucleobase-containing compound | nucleocytoplasmic transpo | 8.681E-07 | | | |
| GO:0006403 | RNA localization | nucleocytoplasmic transpo | 9.775E-11 | | 1.762E-02 | |
| GO:0034504 | protein localization to nucleus | nucleocytoplasmic transpo | 2.303E-02 | | | |
| GO:0071166 | ribonucleoprotein complex localiza | nucleocytoplasmic transpo | 7.783E-10 | | 1.397E-02 | |
| GO:0034502 | protein localization to chromosome | nucleocytoplasmic transpo | 2.440E-02 | | | |
| GO:0051169 | nuclear transport | nucleocytoplasmic transpo | 8.077E-11 | | 7.240E-04 | |
| GO:0007059 | chromosome segregation | chromosome segregation | 2.201E-18 | | | |
| GO:0007017 | microtubule-based process | microtubule-based proces | 2.010E-05 | | | |
| GO:0022402 | cell cycle process | cell cycle process | 2.262E-32 | | | |
| GO:0051304 | chromosome separation | cell cycle process | 1.157E-05 | | | |
| GO:0044770 | cell cycle phase transition | cell cycle process | 1.503E-18 | | | |
| GO:0007051 | spindle organization | cell cycle process | 6.443E-15 | | | |
| GO:0051301 | cell division | cell division | 9.813E-14 | | | |
| GO:0007049 | cell cycle | cell cycle | 3.231E-33 | | 8.364E-04 | |
| GO:0006974 | cellular response to DNA damage | cellular response to DNA | 1.478E-18 | | 2.438E-02 | |
| GO:0072331 | signal transduction by p53 class m | cellular response to DNA | 7.634E-09 | 3.779E-02 | | |
| GO:0010033 | response to organic substance | cellular response to DNA damage stimulus | | | 9.704E-03 | |
| GO:0031668 | cellular response to extracellular st | cellular response to DNA damage stimulus | | | 3.561E-02 | |
| GO:0006950 | response to stress | cellular response to DNA damage stimulus | | | 8.027E-04 | |

| | | | | | | |
|---|---|---|---|---|---|---|
| GO:0035556 | intracellular signal transduction | cellular response to DNA damage stimulus | | | 3.500E-05 | |
| GO:0009408 | response to heat | cellular response to DNA damage stimulus | | | | 4.087E-02 |
| GO:0034620 | cellular response to unfolded prote | cellular response to DNA damage stimulus | | | | 2.720E-02 |
| GO:0042770 | signal transduction in response to | cellular response to DNA | 1.153E-02 | 1.883E-02 | | |
| GO:0000724 | double-strand break repair via hom | cellular response to DNA | 1.557E-05 | | | |
| GO:0000725 | recombinational repair | cellular response to DNA | 2.144E-05 | | | |
| GO:0051716 | cellular response to stimulus | cellular response to DNA damage stimulus | | | 5.722E-06 | |
| GO:0006302 | double-strand break repair | cellular response to DNA | 1.214E-06 | | | |
| GO:0071840 | cellular component organization or | cellular component organi | 1.085E-11 | | 1.412E-02 | |
| GO:0007154 | cell communication | cell communication | | | 1.244E-02 | |
| GO:0070988 | demethylation | demethylation | | | | 3.861E-07 |
| GO:0032259 | methylation | methylation | 3.424E-05 | | | |
| GO:0046034 | ATP metabolic process | ATP metabolic process | | 3.985E-03 | | |
| GO:0006119 | oxidative phosphorylation | oxidative phosphorylation | | 1.601E-03 | | |
| GO:0006123 | mitochondrial electron transport cy | oxidative phosphorylation | | 3.019E-03 | | |
| GO:0022900 | electron transport chain | oxidative phosphorylation | | 2.228E-03 | | |
| GO:1901360 | organic cyclic compound metabolic | organic cyclic compound | 1.154E-45 | | 2.764E-06 | |
| GO:0006807 | nitrogen compound metabolic proc | organic cyclic compound | 2.302E-35 | 9.327E-03 | 3.429E-04 | |
| GO:0009058 | biosynthetic process | organic cyclic compound | 5.392E-19 | 3.398E-04 | | |
| GO:0043170 | macromolecule metabolic process | organic cyclic compound | 9.544E-32 | | | |
| GO:0044237 | cellular metabolic process | organic cyclic compound | 1.209E-32 | 9.450E-04 | 3.639E-04 | |
| GO:0044238 | primary metabolic process | organic cyclic compound | 1.394E-29 | 2.176E-02 | 3.280E-05 | |
| GO:0071704 | organic substance metabolic proce | organic cyclic compound | 2.481E-26 | | | |
| GO:0006091 | generation of precursor metabolite | generation of precursor metabolites a | | 4.334E-02 | | |
| GO:0046483 | heterocycle metabolic process | generation of precursor m | 1.025E-50 | | 1.038E-05 | |
| GO:0006793 | phosphorus metabolic process | generation of precursor metabolites and energy | | | 1.292E-04 | |
| GO:0006725 | cellular aromatic compound metab | generation of precursor m | 5.388E-50 | | 4.120E-06 | |
| GO:0034641 | cellular nitrogen compound metabo | generation of precursor m | 2.593E-44 | 2.099E-08 | 5.171E-04 | |
| GO:1901564 | organonitrogen compound metabo | generation of precursor m | 6.698E-04 | 1.161E-02 | | |
